# Proteomic analysis of spermathecal fluid reveals factors related to long-term sperm storage in ant queens

**DOI:** 10.1101/2022.11.02.513948

**Authors:** Ayako Gotoh

**Affiliations:** Department of Biology, Faculty of Science and Engineering, Konan University, Kobe 658-8501, Japan; Institute for Integrative Neurobiology, Konan University, Kobe 658-8501, Japan; Suntory Rising Stars Encouragement Program in life Sciences (SunRiSE)

**Keywords:** Ant queen, long-term sperm storage, spermatheca, spermathecal fluid, proteomics

## Abstract

Ant queens can maintain a large number of sperm cells for over a decade after mating at the beginning of their adult lives until they die. This ability is prominent because sperm cells cannot maintain their fertilization ability long after ejaculation in animals; however, the cellular mechanisms remain unclear. Sperm cells are maintained in the female sperm storage organ, the spermatheca, which supplies a suitable environment for sperm cells. To reveal the molecular basis of the long-term sperm storage mechanisms in ant queens, protein profiles enriched in the spermathecal fluid relative to the hemolymph were identified in *Lasius japonicus* using mass spectrometry-based proteomics. Proteins related to the extracellular matrix, antioxidant, metabolic pathways, proteases, and with uncharacterized functions were enriched in the spermathecal fluid relative to the hemolymph. These enriched proteins were shared with highly expressed genes previously detected by transcriptome analyses of the spermatheca in queens of *Crematogaster osakensis* belonging to a different subfamily than *L. japonicus*. It is considered that the ability for long-term sperm storage has evolved in the early ant lineage; therefore, the common proteins identified in the two ant species are crucial for this ability.

## 1. Introduction

Queens of ants and honeybees, which develop prominent high eusociality, show remarkably high reproductive abilities, including not only large amounts of egg production but also prolonged sperm storage. In contrast to other insect females that mate repeatedly throughout their life, queens of social Hymenoptera, such as ants, wasps, and bees mate only for a short period in their early adult lives and produce fertilized eggs until the end of their lives without additional mating. The species of paper wasps and hornets distributed in the temperate zone live approximately a year (Gotoh et al., 2008), while the honey bee *Apis mellifera* lives for approximately 2–4 years (Dade, 1994). Queen in many ant species can live for more than a decade (Keller, 1998), indicating that the ant queen is a champion that can store viable sperm for the longest lifespan among animals. This is a unique reproductive ability because sperm cells usually die or lose fertilization ability soon after ejaculation from male animals; however, the mechanisms are still unclear.

Sperm cells are preserved inside the spermathecal reservoir, which has a sac-like structure, soon after mating to before fertilization. Therefore, the spermathecal reservoir contents provide an environment suitable for sperm cells. The columnar epithelial cells localized near the entrance of the spermathecal reservoir presumably have a transporting function but not a secretory function in ant queens (Wheeler and Krutzsch, 1994), and proteins produced in the spermathecal gland, which attaches the spermathecal duct near the opening of the spermathecal reservoir, are secreted into the spermathecal reservoir because gel profiles of proteins are similar between the spermathecal gland and the spermathecal fluid in honeybee queens (Baer et al., 2009; den Boer et al., 2009).

A previous study revealed that stored sperm cells are immobilized and the near-anoxic conditions in the spermathecal reservoir are very important for maintaining sperm immobilization and viability; however, other factors are also necessary for long-term sperm storage in ants (Gotoh and Furukawa, 2018; Gotoh et al., 2022). To clarify the whole picture of long-term sperm storage mechanisms in ant queens, other factors present in the spermathecal fluid should be screened by multi-omics. The first step of this research focused on the protein profiles of spermathecal fluid. Previously, highly expressed genes in the spermatheca relative to the whole body sample were investigated using RNA-seq analyses, and subsequently, genes expressed in the spermathecal gland were identified using the *in situ* hybridization method in ants, *Crematogaster osakensis* (Gotoh et al., 2017). However, it is not clear whether the proteins translated from these genes are secreted into the spermathecal reservoir. Therefore, proteomic analysis of spermathecal fluid is necessary.

To date, proteomic analyses of the spermatheca in leaf-cutting ants, *Atta sexdens rubropilosa* (Malta et al., 2014) and spermathecal fluid in honeybees, *A. mellifera* (Baer et al., 2009) have been performed to understand sperm storage mechanisms. However, these studies only compared the protein profiles of mated and virgin spermathecae. In contrast to other insect females that mate repeatedly throughout their lives, the period available for mating of ants is limited only at the beginning of their adult lives, with no chance to mate after this time. Furthermore, mating dates in ants are unpredictable because they are usually determined by the weather and humidity (Hölldobler and Wilson, 1990). Therefore, it is likely that the environment for the acceptance of sperm cells from males is already established in the spermathecal fluid soon before the mating dates. To date, important candidate genes and proteins for spermathecal functions, such as spermatheca-specific genes in ants and heat shock proteins for temperature tolerance in honeybees, have been screened by comparing the spermatheca and other tissues, but not virgin and mated spermatheca (Gotoh et al., 2017; McAfee et al., 2020). Thus, to characterize the functions of the spermathecal fluid, a comparison between spermatheca and non-spermatheca samples, in which the spermathecal fluid corresponds to the hemolymph, should be appropriate. Moreover, the use of the spermathecal fluid of virgin queens is advantageous because contamination from the proteins of sperm cells can be eliminated. In previous proteomic analyses of the leaf-cutting ants, the protein profiles were investigated from homogenized spermatheca, which includes both proteins from the spermathecal fluid and the epithelial cells of the spermathecal reservoir (Malta et al., 2014); therefore, extraction of only the spermathecal fluid is necessary to identify proteins directly surrounding sperm cells. Furthermore, gel-based proteomic analysis has been the mainstream approach for revealing protein profiles; however, another method, mass spectrometry (MS)-based proteomic analysis, which can identify more protein repertories and determine quantities, has been recently developed (Xu et al., 2021). This study identified proteins enriched in the spermathecal fluid relative to the hemolymph of *Lasius japonica* queens using data-independent acquisition (DIA)-based quantitative proteomics technology, which is one of the MS-based proteomics analyses to reveal molecules affecting long-term sperm storage in ants.

## 2. Material and Methods

### 2.1. Ants

*L. japonica* colonies containing alate virgin queens were collected before nuptial flight in July 2021 at Kobe City in Hyogo Prefecture. The ants were housed in plastic containers with moistened plaster bases. This ant species is useful for investigating spermathecal fluid because their spermathecal reservoir size is large among ant species.

### 2.2. Sampling of spermathecal fluid and hemolymph

Alate queens were dissected without anesthesia to avoid changes in protein expression, and their spermathecae were placed in 80 *μ*L distilled water for high-performance liquid chromatography (HPLC; Fujifilm Wako Pure Chemical Corporation, Japan). Subsequently, they were washed six times to avoid contamination of hemolymph and fat bodies. After complete removal of the water around the spermatheca, a hole was created in the spermathecal reservoir using a tungsten needle. Because the columnar epithelium in the hilar region of the spermathecal reservoir contains enzymes related to energy metabolism due to the presence of abundant mitochondria (Wheeler and Krutzsch, 1994), this region was avoided when creating the hole to prevent contamination of the cells. The spermathecal fluid was then collected via the small hole by inserting a fine glass capillary and was subsequently released into 5 *μ*L distilled water for HPLC in a microtube on ice. Spermathecal fluid collected from 23–25 queens was pooled into one microtube and frozen at −80 °C. Prior to proteomic analysis, spermathecal fluids in two microtubes were combined into one microtube as one sample. For hemolymph sampling, a hole was created between the segment of the side prothorax and the mesothorax using a tungsten needle, and then the hemolymph was collected using a fine glass capillary without contamination of the fat body. The hemolymph collected from 10 queens was ejected into 5 *μ*L distilled water for HPLC in a microtube on ice and frozen at −80 degree. The samples on dry ice were then shipped to Kazusa Genome Technologies (Japan) for proteomic analyses.

### 2.3. Proteomic sample preparation

Acetonitrile (ACN) containing 0.1% trifluoroacetic acid (TFA) was added to the sample to inactivate the proteins, which were then dried using a centrifugal evaporator. The dried sample was extracted in 0.5% sodium dodecanoate and 100 mM Tris-HCl (pH 8.5) using a water-bath-type sonicator (Bioruptor II; Cosmo Bio Inc., Tokyo, Japan) for 15 min. The protein extract was treated with 10 mM dithiothreitol at 50 °C for 30 min and alkylated with 30 mM iodoacetamide in the dark at room temperature for 30 min. The iodoacetamide reaction was stopped by treatment with 60 mM cysteine for 10 min. The mixture was diluted with 150 *μ*L of 50 mM Tris-HCl pH8.0 and digested by adding 500 ng Trypsin/Lys-C mix (Promega, Madison, WI, USA) overnight at 37 °C. The digested sample was acidified with 30 *μ*L of 5% TFA, followed by sonication (Bioruptor UCD-200) for 5 min. The mixture was shaken for 5 min and centrifuged at 15,000 g for 5 min. The supernatant was desalted using C18-StageTips and dried using a centrifugal evaporator. Dried peptides were dissolved in 3% ACN and 0.1% TFA. The redissolved peptides were measured using a colorimetric peptide assay kit (Thermo Fisher Scientific, MA, USA) and transferred to a hydrophilic-coated low adsorption vial (ProteoSave vial; AMR Inc., Tokyo, Japan).

### 2.4. MS Analysis

Peptides (500 ng) were directly injected onto an Aurora Series emitter column (25 cm × 75 μm ID, 1.6 μm, C18; IonOpticks, Melbourne, Australia) at 60 °C and then separated with a 120 min gradient at a flow rate of 200 nl/min using an UltiMate 3000 RSLCnano system (Thermo Fisher Scientific). Peptides eluted from the column were analyzed using an Orbitrap Exploris 480 mass spectrometer (Thermo Fisher Scientific) for overlapping window DIA-MS (Amodei et al., 2019; Kawashima et al., 2019). MS1 spectra were collected in the range of 495–745 m/z at 15,000 resolution with an automatic gain control (AGC) target of 3e6 and the maximum injection time set at “auto”. MS2 spectra were collected at >200 m/z and 45,000 resolution with an AGC target of 3e6, maximum injection time set to “auto,” and stepped normalized collision energies of 22, 26, and 30%. The isolation width for MS2 was set to 4 m/z and overlapping window patterns at 500– 740 m/z were used as window placements optimized by Skyline v4.1 (MacLean et al., 2010).

### 2.5. Proteomic data analysis

MS files were screened against a *Lasius niger* library using Scaffold DIA (Proteome Software Inc., Portland, OR, USA). The *L. niger* spectral library was generated from the *L. niger* UniProtKB/Swiss-Prot protein sequence database (UniProt ID UP000036403, 18140 entries) using Prosit (Gessulat et al., 2019; Searle et al., 2020). The scaffold DIA search parameters were as follows: experimental data search enzyme, trypsin; maximum missed cleavage sites, 1; precursor mass tolerance, 10 ppm; fragment mass tolerance, 10 ppm; static modification, cysteine carbamidomethylation; and q-value, < 0.1. The protein identification threshold was set at both peptide and protein false discovery rates of less than 1%. Peptide quantification was performed using the EncyclopeDIA algorithm (Gessulat et al., 2019) in Scaffold DIA. For each peptide, the five highest-quality fragment ions were selected for quantitation. Protein quantification was performed using summed peptide quantification. The protein quantification data were transformed to log2 (protein intensities) and filtered so that at least one group contained a minimum of 70% valid values for each protein. The remaining missing values were imputed using random numbers drawn from a normal distribution (width, 0.3; downshift, 1.8) in Perseus v1.6.15.0 (Tyanova et al., 2016). The thresholds for altered proteins showed more than two-fold change (Welch test, p < 0.05), which differed between the two groups.

### 2.6. Integration of the spermatheca data from a previous study into RNA-sequencing data

To integrate enriched proteins detected by proteomic analysis in this study and by transcriptomic analysis in a previous study (Gotoh et al., 2017), BlastP homology search of the amino acid sequences of the proteins detected in this study was performed against amino acid sequences originating from highly expressed genes of the spermatheca relative to the whole body at 1 week and 1 year after mating (cut-off e-value=1e-100). When multiple *C. osakensis* sequences were aligned to a *L. niger* protein, the *C. osakensis* sequences with the smallest evalue score corresponded.

### 2.7. Gene ontology and pathway enrichment analysis

For gene ontology (GO) and Kyoto Encyclopedia of Genes and Genomes (KEGG) pathway analyses, proteins of *L. niger* were converted to orthologous proteins of the honeybee, *Apis mellifera*, which is known as a model organism of social Hymenoptera, because information on gene annotation and pathway analysis are well developed in honeybees. Amino acid sequences of *L. niger* and *A. mellifera* proteins were obtained from the Uniprot database (https://www.uniprot.org) (The UniProt, 2021), and a BlastP homology search of the amino acid sequences of 3069 proteins detected in this study was performed against the sequences of 21802 proteins in *A. mellifera* (cut-off e-value=1e-50). When multiple *A. mellifera* proteins were aligned to a *L. niger* protein, the *A. mellifera* protein with the smallest e-value score corresponded.

GO enrichment analysis was performed using ShinyGo 0.76.2 (http://bioinformatics.sdstate.edu/go/) (Ge et al., 2020) with default parameters by inputting UniProt ID of proteins enriched in the spermathecal fluid and hemolymph. Of the 3069 proteins, 2770 showed similarity to *A. mellifera* proteins (1578 of 1707 and 157 of 202 proteins enriched in spermathecal fluid and hemolymph, respectively). For KEGG enrichment analysis, *A. mellifera* Uniprot IDs were converted to Entrez IDs using ID mapping hosted by UniProt (https://www.uniprot.org/id-mapping). Of the 2770 *A. mellifera* Uniprot IDs, 1735 were converted to Entrez IDs (1012 of 1707 and 78 of 202 *L. niger* proteins enriched in spermathecal fluid and hemolymph, respectively). KEGG pathways with relative protein expression levels between spermathecal fluid and hemolymph were visualized using Pathview (https://pathview.uncc.edu) (Luo et al., 2017) by inputting Entrez IDs and values of Log2 fold changes.

## 3. Results and Discussion

### 3.1. Overview

A total of 3069 proteins were detected in this analysis. The number of identified proteins is 1–2 order of magnitude greater than in previous proteome studies of the spermatheca in ants and honeybees (Baer et al., 2009; Malta et al., 2014). Therefore, the protein profiling of the spermathecal fluid should be deeply characterized at high resolution by MS analyses. Among the 3069 proteins detected, 1707 and 202 proteins were enriched in the spermathecal fluid and hemolymph, respectively (*P*-value <0.05, llog_2_ fold changel>1, Fig. 1). The number of proteins enriched in the spermathecal fluid was approximately 8.5 times more than those in the hemolymph, suggesting that various proteins with various functions that form complicated molecular networks are essential for sperm maintenance.

**Fig. 1.**
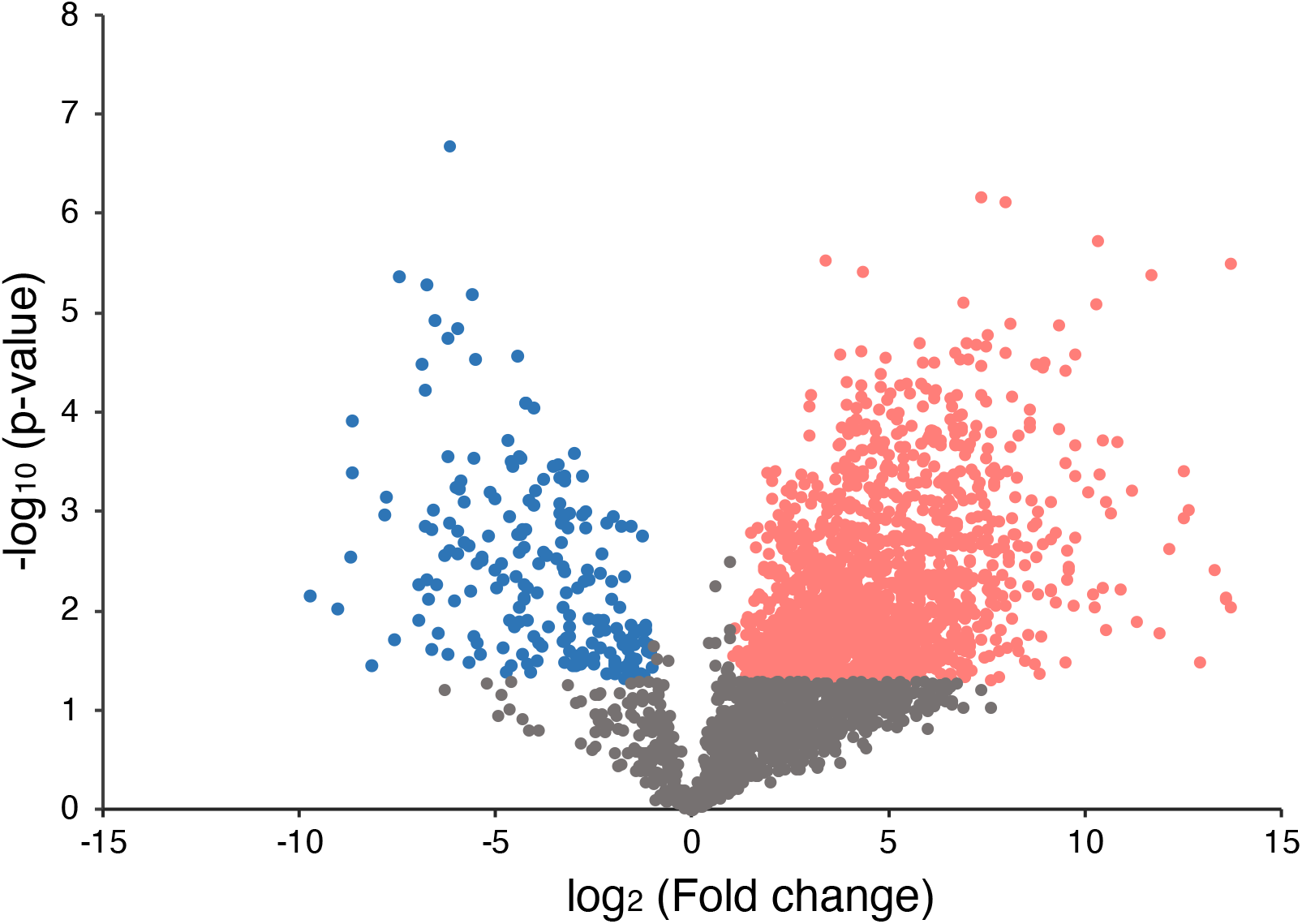
Volcano plot representing differences in protein expression between the spermathecal fluid and hemolymph. Significantly enriched proteins in the spermathecal fluid and hemolymph are shown in pink and blue, respectively (>2 fold or < 2 fold, p < 0.05).

These proteins enriched in the spermathecal fluid were categorized into spermatheca-specific genes and proteins related to the extracellular matrix, antioxidant, metabolic process, protease, and chaperone, as explained in the following sections.

### 3.2. Proteins identical proteins of spermatheca-specific genes detected by transcriptomic analysis

Of the 1707 proteins enriched in the spermathecal fluid, 142 proteins were homologous to the proteins originating from highly expressed genes detected in the spermatheca relative to the whole body at both 1 week and 1 year after mating in queens of *C. osakensis* (Fig. 2, Table 1, and Table S1; Gotoh et al., 2017).

**Fig. 2.**
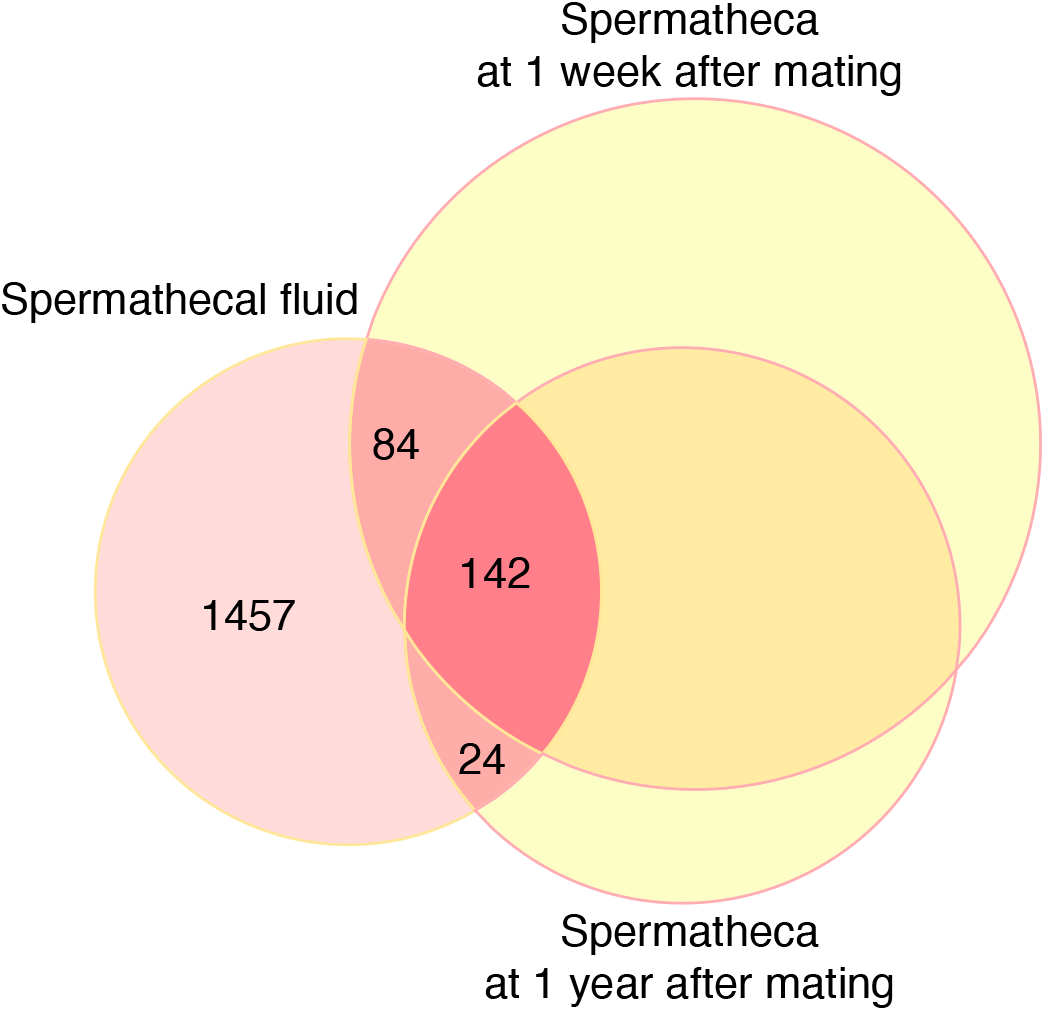
Venn diagram showing the number of common enriched genes and proteins in the spermatheca 1 week and 1 year after mating relative to the body samples of *Crematogaster osakensis* queens (Data from Gotoh et al. 2017) and in the spermathecal fluid relative to the hemolymph in *Lasius japonicus* queens (Data from this study).

**Table 1.** List of selected highly localized proteins in the spermathecal fluid relative to hemolymph.

| Proteome data |  | Highly expressed genes in the spermatheca relative to the body in *C. osakerensis* (Gotoh et al., 2017). |  |  |  |  |  |
| --- | --- | --- | --- | --- | --- | --- | --- |
| Category | Uniprot Accession | Protein Name | Log2FC | Contig. No. | Log2FC (1 week) | Log2FC (1 year) | Expression pattern* |
| Spermatheca specific genes | A0A0J7KLS0 | Collagen alpha-1 chain | 9.8 | comp63322_c0_seq1.m.59242 | 8.7 | 9.6 | SG |
|  | A0A0J7P5F7 | ZP domain-containing protein | 7.3 | comp65548_c0_seq1.m.79879 | 8.8 | 8.5 | HE |
|  | A0A0J7KQ48 | Protein unc-13-like protein | 4.6 | comp65714_c0_seq1.m.81456 | 2.7 | 2.6 | SG |
| Antioxidant | A0A0J7NA12 | Catalase (Fragment) | 5.4 | - | - | - | - |
| " | A0A0J7K2I5 | Glutathione S-transferase (Fragment) | 5.1 | - | - | - | - |
| " | A0A0J7K579 | Thioredoxin domain-containing protein 4 | 4.6 | comp70443_c0_seq1.m.279284 | 1.0 | Not significant | - |
| " | A0A0J7KSQ9 | Superoxide dismutase | 4.4 | comp65723_c0_seq1.m.35720 | 1.1 | 1.3 | HE, MG, OV |
| " | A0A0J7KUV3 | Chorion peroxidase | 4.1 | comp70976_c0_seq1.m.342323 | 6.3 | 5.1 | Failed |
| " | A0A0J7L5D0 | Glutathione synthase | 3.7 | comp69937_c0_seq12.m.230072 | 2.5 | Not significant | - |
| Chaperone | A0A0J7P4F1 | DnaJ-like subfamily c member 3-like protein | 6.8 | - | - | - | - |
| " | A0A0J7L4V4 | Heat shock protein 83 | 3.7 | - | - | - | - |
| " | A0A0J7KVR3 | Heat shock 70 kDa protein cognate 5 | 3.6 | comp69851_c2_seq1.m.223848 | 2.3 | 1.7 | HE, MG, OV, SG |
| " | A0A0J7K484 | Hypoxia up-regulated protein 1 | 3.7 | - | - | - | - |
| " | A0A0J7KBH0 | Hypoxia up-regulated protein 1 | 2.9 | - | - | - | - |
| Extracellular matrix related | A0A0J7L2K3 | Cuticlin-1 | 13.6 | - | - | - | - |
| " | A0A0J7KHK2 | Papilin | 10.3 | comp67808_c0_seq1.m.123784 | 2.9 | 1.5 | - |
| " | A0A0J7KYI3 | Protein slit (Fragment) | 6.9 | comp65666_c1_seq1.m.81012 | 3.9 | 2.6 | - |
| " | A0A0J7KRL3 | Chondrotin proteoglycan-2-like protein | 5.6 | comp63194_c1_seq1.m.58387 | 6.7 | 5.1 | HE, MG, OV, SG |
| " | A0A0J7LOV3 | Thrombospondin type-1 domain-containing protein 4 | 4.7 | comp66057_c1_seq1.m.86327 | 4.5 | 4.2 | CO, FB, GC, HE, MG, OV, SD, SG |
| " | A0A0J7L7A2 | Procollagen-oxoglutarate 5-dioxygenase 3 | 3.3 | comp61774_c0_seq1.m.51004 | Not significant | 1.0 | - |
| " | A0A0J7L8E5 | Peroxidase-like protein | 2.4 | comp70976_c0_seq1.m.342323 | 6.3 | 5.1 | Failed |
| Protease | A0A0J7LAQ5 | Serine protease inhibitor dipetabagastin isoform x3 | 13.3 | - | - | - | - |
| " | A0A0J7KNW4 | Carboxypeptidase b-like protein | 12.9 | - | - | - | - |
| " | A0A0J7KPY2 | Serine proteinase stubble | 12.5 | - | - | - | - |
| " | A0A0J7NWH4 | Serine proteinase stubble | 11.2 | - | - | - | - |
| " | A0A0J7LOF7 | Trypsin 3a1 | 10.4 | - | - | - | - |
| " | A0A0J7NF30 | Serine proteinase stubble | 7.1 | - | - | - | - |
| Energy metabolism | A0A0J7NMH8 | Glucose dehydrogenase | 7.6 | comp71645_c0_seq1.m.442441 | 3.0 | 3.4 | MG, SG |
| " | A0A0J7MY75 | Succinate-CoA ligase [GDP-forming] subunit beta, mitochondrial | 4.3 | comp63688_c0_seq1.m.34877 | 1.9 | 1.5 | - |
| " | A0A0J7LAF5 | L-lactate dehydrogenase | 3.1 | comp55829_c0_seq1.m.33328 | 3.7 | 3.9 | - |
| " | A0A0J7NNS3 | Malate dehydrogenase | 3.1 | comp70443_c1_seq3.m.279322 | 1.4 | Not significant | - |
| " | A0A0J7KQB8 | Glucose-6-phosphate 1-dehydrogenase | 2.6 | comp71903_c0_seq1.m.489095 | 2.2 | 1.6 | - |
| " | A0A0J7KNC5 | Glutamate dehydrogenase (NAD(P)(+)) (Fragment) | 2.3 | comp70263_c4_seq1.m.260523 | 3.1 | 2.5 | HE, MG, OV, SG |
| Others | A0A0J7KXL2 | Isoform x (Fragment) | 13.7 | comp71814_c1_seq10.m.473823 | 1.7 | 1.8 | - |
| " | A0A0J7JV67 | Xanthine dehydrogenase (Fragment) | 6.5 | - | - | - | - |
| " | A0A0J7KKD4 | Ejaculatory bulb-specific protein 3 (Fragment) | 3.7 | - | - | - | - |
| " | A0A0J7L6X2 | Bee-milk protein | 10.5 | - | - | - | - |
| " | A0A0J7KDK5 | Uncharacterized protein | 10.2 | comp65472_c0_seq1.m.78985 | 2.3 | 1.7 | - |
| " | A0A0J7NL95 | Uncharacterized protein | 9.8 | comp71363_c1_seq1.m.401420 | 2.8 | 2.6 | - |
| " | A0A0J7LOG4 | Uncharacterized protein | 6.1 | comp71609_c2_seq1.m.435392 | 1.6 | 1.0 | - |
| " | A0A0J7KW64 | Uncharacterized protein | 6.0 | comp66952_c1_seq1.m.103070 | 3.2 | 2.6 | FB, GC, HE, MG, OV, SD, SG |

\* Failed = no signals, SG = spermathecal gland, HE = hilar columnar epithelial cells of spermatheca reservoir, SD = spermathecal duct, OV = ovary, MG = midgut, CO = common oviduct, FB = fat body

Three enriched proteins, unc-13-like protein (A0A0J7KQ48), collagen alpha-1 chain (A0A0J7KLS0), and ZP domain-containing protein (A0A0J7P5F7), were homologous to the amino acid sequences of spermatheca-specific genes, as revealed by transcriptome and *in situ* hybridization analyses (Gotoh et al., 2017). The former two genes were expressed only in the spermathecal gland, suggesting that these proteins were translated and secreted into the spermathecal reservoir, while the latter gene was expressed only in the columnar epithelium of the spermathecal reservoir, suggesting that these transporters regulate the passage of this protein after translation because of the absence of secretory function (Wheeler and Krutzsch, 1994). The UNC-13 protein is necessary for synaptic vesicle exocytosis during neurotransmission in *Drosophila melanogaster* and *Caenorhabditis elegans* (Aravamudan et al., 1999; Richmond et al., 1999). However, its functions in the reproductive tract have not been reported; therefore, this protein has novel functions in sperm storage in ants.

Xanthine dehydrogenase (A0A0J7JV67) was also abundant in the spermathecal fluid, and this gene is also spermathecal gland-specific; however, its amino acid sequences are not homologous (Table 1). The reason for this is unclear, but they may be isoforms that may have different functions in sperm storage.

Because long-term sperm storage by ant queens is a common trait among many ant species belonging to various subfamilies, the mechanisms of prolonged sperm storage should have evolved in the early ant lineage. *C. osakensis* used in transcriptome analysis and *Lasius* species in this study belong to different subfamilies in the family Formicidae (Brady et al., 2006). Therefore, common proteins enriched in the spermathecal fluid and spermatheca are crucial candidates for long-term sperm storage.

### 3.3. Extracellular matrix-related proteins

Including the two proteins mentioned in section 3.2. that are homologous to spermatheca-specific genes, enrichment of many extracellular matrix-related proteins was conspicuously found both in the spermathecal fluid and spermatheca (Table 1). However, in a previous histological study, collagen fiber-like ultrastructures were never reported (Wheeler and Krutzsch, 1994). These may play a role in physical protection by providing viscosity to the spermathecal fluid. Their localization and function remain unknown; therefore, further research is required.

### 3.4. Antioxidant enzymes

Antioxidant enzymes such as glutathione S-transferase, catalase, and superoxide dismutase were enriched in the spermathecal fluid (Table 1). A part of the glutathione metabolism pathway also tended to be activated by proteins enriched in the spermathecal fluid (Fig. S1). Furthermore, GO enrichment analyses revealed that molecular functions related to oxidoreductase activity were enhanced in the spermathecal fluid relative to the hemolymph (Fig. 3). Reactive oxygen species (ROS) cause damage to sperm cellular functions, and sperm DNA as well as various antioxidants are found in the semen of mammals (Aitken, 2017; Sanocka and Kurpisz, 2004). The expression and localization of antioxidant enzymes have also been reported in the spermathecae of honey bees (Baer et al., 2009; Collins et al., 2004; Gonzalez et al., 2018; Rangel et al., 2021; Weirich et al., 2002) and ants (Gotoh et al., 2017). Furthermore, the interior of the spermatheca is a near-anoxic environment in honeybees and ants (Gotoh et al., 2022; Paynter et al., 2017). Double strategies against ROS should be crucial for maintaining sperm fertilization ability over the long term.

**Fig. 3.**
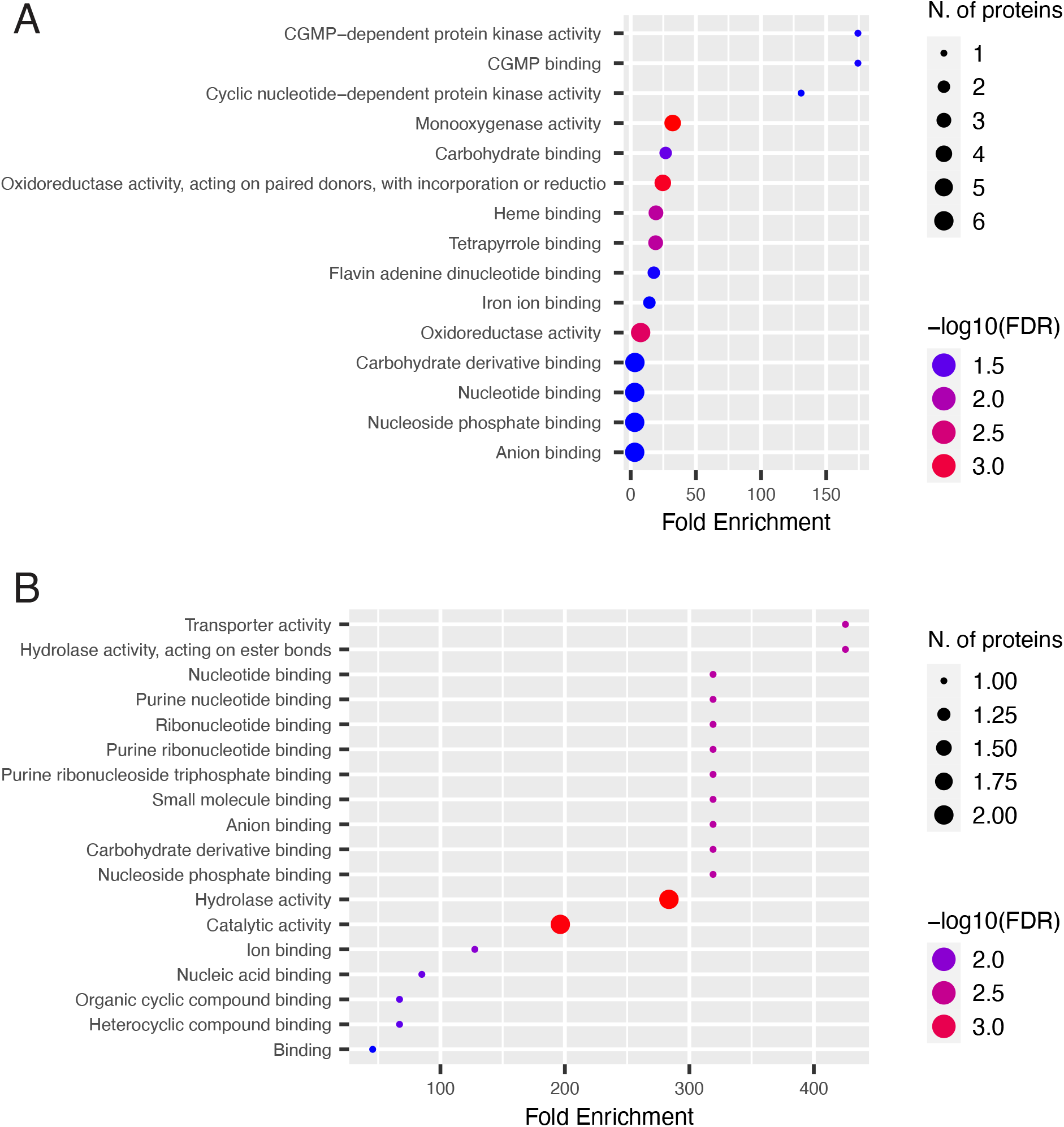
Gene ontology (GO) enrichment analysis by molecular function for enriched proteins in the spermathecal fluid (A) and hemolymph (B) ranked by fold enrichment values. The size and colors of the circle represent the number of proteins in the GO classification and the −log10 of the false-discovery rate (FDR), respectively.

### 3.5. Proteins related to metabolic processes

Enriched proteins related to energy production were found in the spermathecal fluid as along with genes enriched in the spermatheca (Table 1; Gotoh et al., 2017). This indicates that the enzymes are secreted into the spermathecal fluid and may contribute to the maintenance of sperm metabolism. Enhancement of the tricarboxylic acid (TCA) cycle was more conspicuous than that of the glycolytic pathway in the spermathecal fluid relative to the hemolymph by KEGG pathway analyses (Fig. 4). This is not consistent with spermathecal fluid analyses of honeybee queens, where the glycolytic pathway gives precedence to sperm storage in a hypoxic spermathecal environment (Baer et al., 2009; Paynter et al., 2017). This difference may be due to the different traits of each species or the use of different proteomic methods. The ability of queens to store sperm long-term has convergently evolved in ants and honeybees belonging to different hymenopteran lineages. Therefore, it is intriguing whether differences exist in prolonged sperm storage mechanisms.

**Fig. 4.**
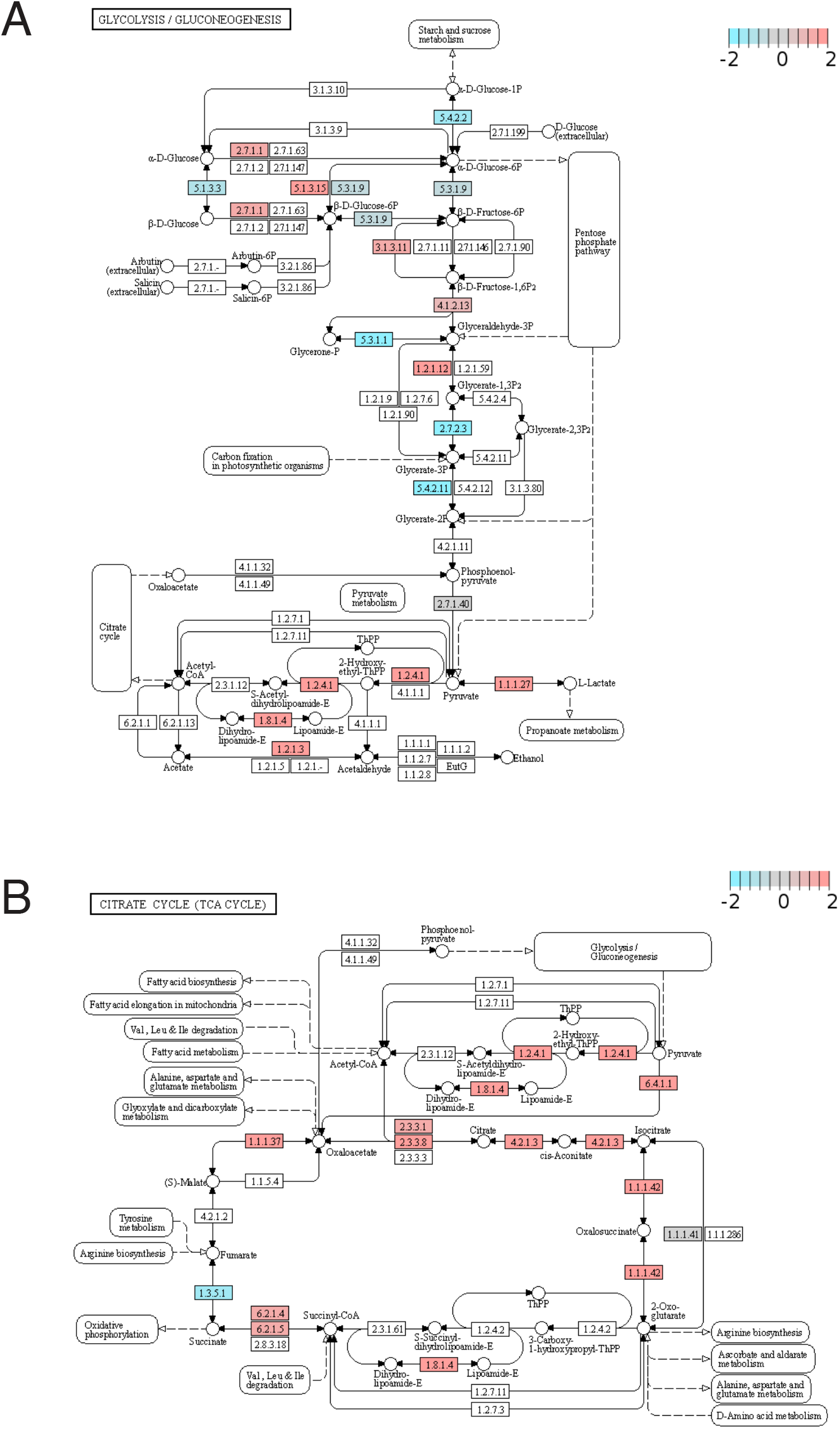
The KEGG pathway of glycolysis (A) and TCA cycle (B). The colored boxes (pink, grey, and blue) correspond to the proteins detected in this study. The color intensity indicates expression levels of proteins enriched in the spermathecal fluid (pink) and hemolymph (blue).

In amino acid metabolism, spermathecal fluid proteins were abundantly assigned to the degradation pathways of valine, leucine, and isoleucine (Fig. S2A) but not in the biosynthesis pathway (Fig. S2B). These degradation pathways may play a role in the elimination of these amino acids released from sperm cells.

The inositol phosphate pathway also tended to be activated in the spermathecal fluid (Fig. S3). Inositol phosphate is known to play an important role in cell signaling; however, its importance in sperm storage has not been investigated.

Further metabolomic analyses of the spermathecal fluid and sperm cells are necessary to reveal how metabolites are exchanged and processed between the spermathecal fluid and sperm cells.

### 3.6. Proteases

Proteases, such as trypsin, were especially enriched in the spermathecal fluid with very high fold changes (Table 1), with three and five proteases ranked in the top 10 and 20 enriched proteins, respectively (Table S1). This trend is consistent with the transcriptome data of the spermatheca; however, their amino acid sequences were not homologous (Table 1; Gotoh et al., 2017). Proteases have various functions associated with reproduction, such as sperm motility in males of lepidopteran species and water striders (Friedländer et al., 2001; Miyata et al., 2012; Osanai et al., 1989) as well as maturation in mammals (Cesari et al., 2010). Recently, serine proteases from seminal fluid were reported to decrease sperm viability of rival males as a result of sperm competition, but serine protease inhibitors from spermathecal fluid prevented sperm competition in *Atta colombica* ants (Dosselli et al., 2019). Reproduction-related proteins are thought to evolve rapidly under positive selection (Findlay and Swanson, 2010; Prokupek et al., 2010). In fact, serine proteases expressed in female reproductive organs have evolved under positive selection in the mosquito, *Anopheles gambiae* (Mancini et al., 2011). The inconsistency in the amino acid sequences of the proteases may be due to rapid molecular evolution. It could also be considered that these proteases play a role in different phases, where the proteases detected here may be related to male-female conflict during or soon after copulation because the spermathecal fluid was sampled from queens soon before mating, whereas the proteases detected by transcriptome analysis may be associated with long-term sperm maintenance because the expression levels remain high in the spermatheca of queens at both 1 week and 1 year after mating (Gotoh et al., 2017).

### 3.7. Chaperones

Proteins that matched chaperones, including heat shock proteins, were also identified (Table 1). Protein chaperone is known to have a role in prevention of protein aggregation and misfolding (Fink, 1999). These proteins may maintain the function of sperm or spermathecal fluid. Hypoxia upregulated protein 1 (A0A0J7K484 and A0A0J7KBH0) was identified in the proteome analyses, but not in the transcriptome analysis (Table 1). This protein belongs to the heat shock protein 70 family and is involved in protein folding (Wang et al., 2021). This function may be related to sperm protection in a near-anoxic environment in the spermathecal reservoir (Gotoh et al., 2022).

### 3.8. Conclusion

This study narrowed down a list of candidate proteins that likely play important roles in long-term sperm storage in ant queens, including not only antioxidants and chaperones, which have been reported frequently in honeybee studies, but also unc-13-like proteins and extracellular matrix-related proteins, which have not been studied previously. This was accomplished because 1) the protein profiles of the spermathecal fluid and hemolymph were compared, but those of mated and virgin spermathecal were not, 2) a large number of proteins were identified and quantified at high resolution using MS-based proteomic technology, and 3) the protein components of the spermathecal fluid were compared to the results of a previous transcriptomics study of the spermatheca of *C. osakensis*, which belongs to another ant subfamily.

Further multi-omics analyses, such as metabolomics and lipidomics, and subsequent experiments using molecular techniques such as genome editing and sperm culture with candidate molecules, should provide an understanding of the integrated mechanisms of longterm sperm storage in ant queens.

## Acknowledgments

This work is supported by Grants-in-Aid for Scientific Research from the Japan Society for the Promotion of Science (JSPS) (KAKENHI Grant Numbers: 20K06080) and Suntory Rising Stars Encouragement Program in life Sciences (SunRiSE) to A.G.

## Supplementary Figures

**Fig. S1.**
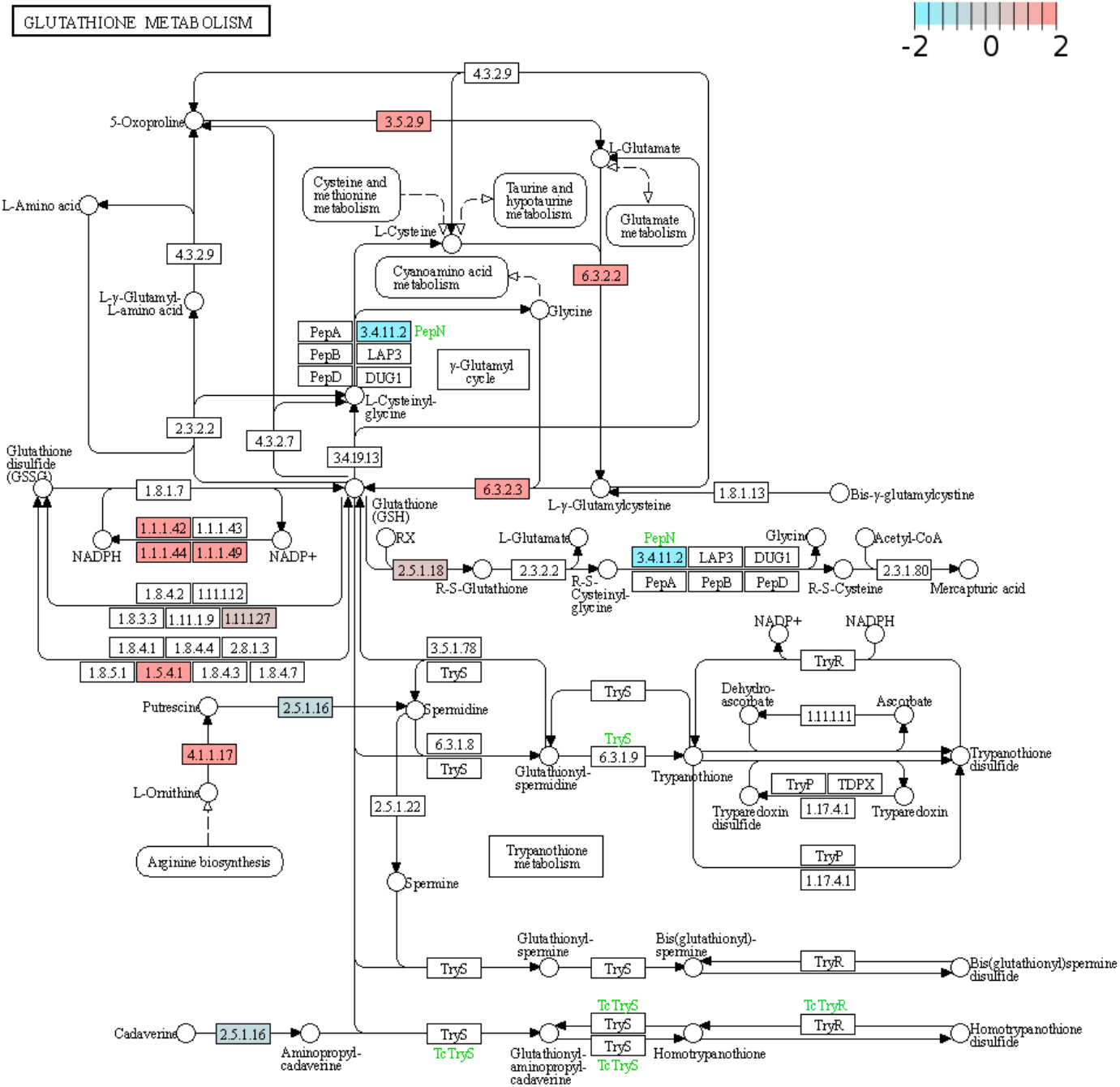
The KEGG pathway of glutathione metabolism. The colored boxes (pink, grey and blue) are corresponding to the proteins detected in this study. The color intensity indicates expression levels of proteins enriched in the spermathecal fluid (pink) and in the hemolymph (blue).

**Fig. S2.**
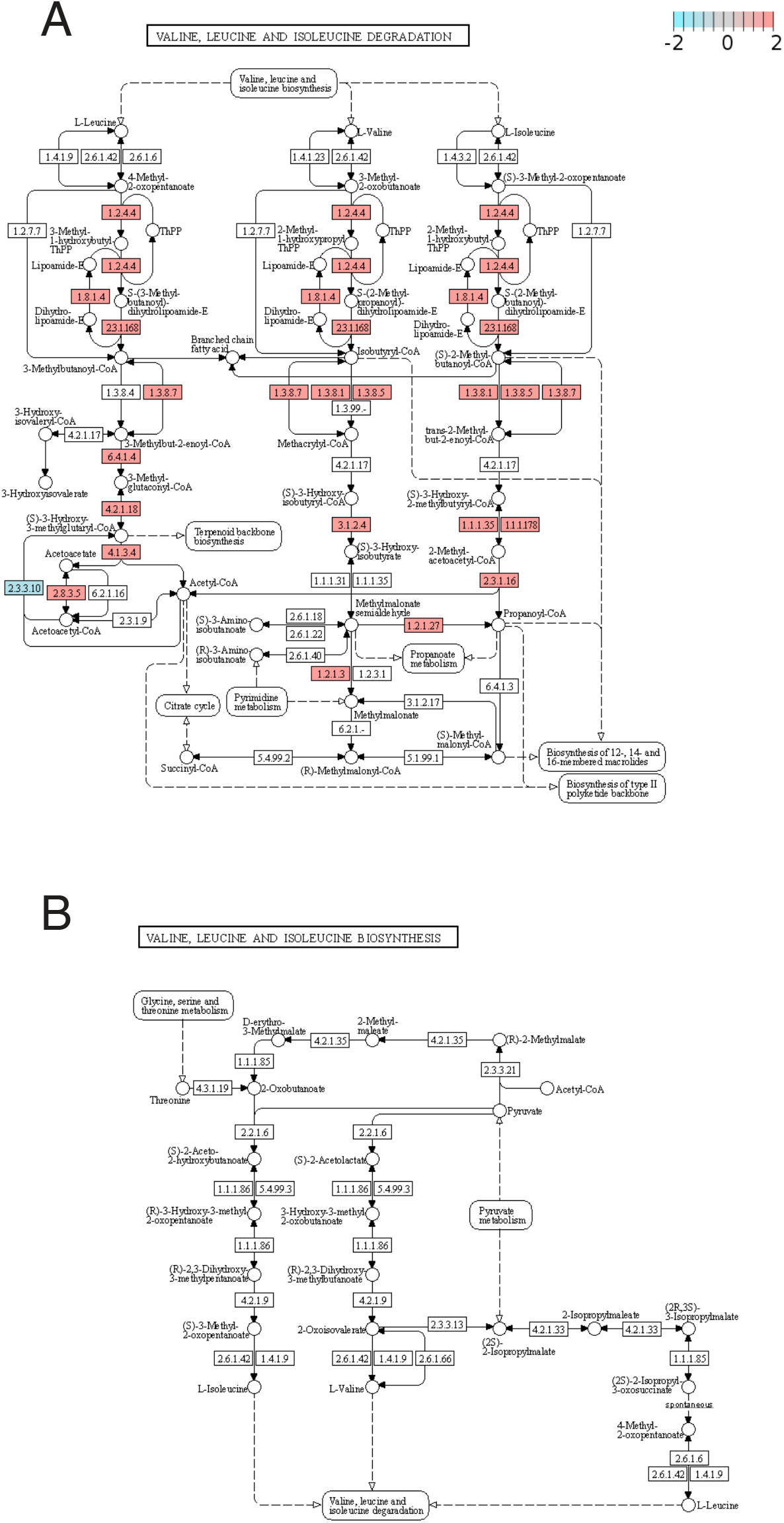
The KEGG pathway of degradation (A) and biosynthesis (B) of valine, leucine and isoleucine. The colored boxes (pink, grey and blue) are corresponding to the proteins detected in this study. The color intensity indicates expression levels of proteins enriched in the spermathecal fluid (pink) and in the hemolymph (blue).

**Fig. S3.**
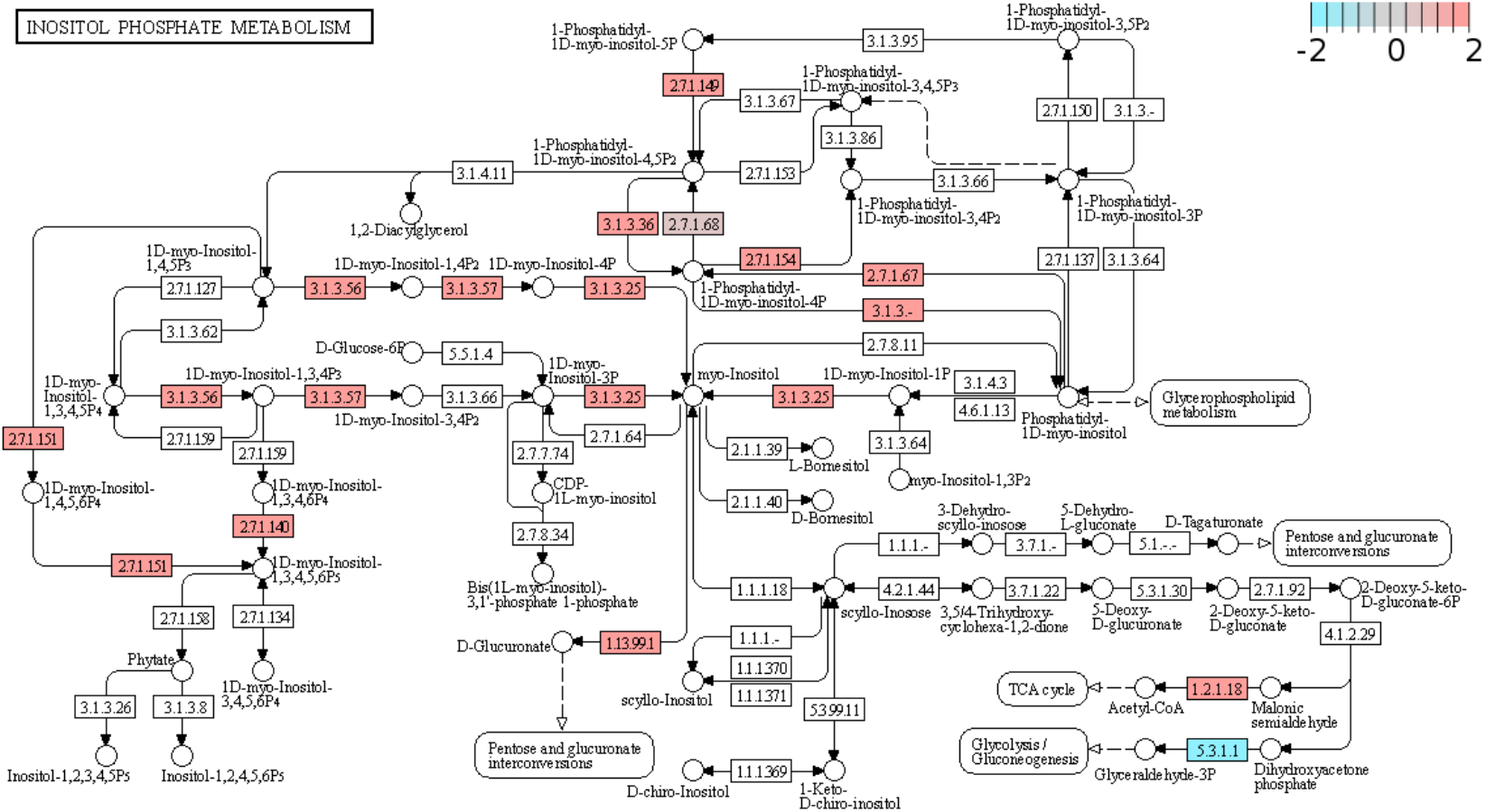
The KEGG pathway of inositol phosphate. The colored boxes (pink, grey and blue) are corresponding to the proteins detected in this study. The color intensity indicates expression levels of proteins enriched in the spermathecal fluid (pink) and in the hemolymph (blue).

